# An immunocompetent osteoblastic model of mammary cancer bone metastasis established by syngeneic intratibial injection of PyMT mammary carcinoma cells in FVB/N mice

**DOI:** 10.64898/2026.07.08.737245

**Authors:** Courtney L. Flatt, Sarah Nano, Ria Goyal, Susan E. Waltz, Glen L. Niebur, Laurie E. Littlepage

## Abstract

Osteoblastic bone metastasis, in which disseminated tumor cells drive net bone formation, is a clinically distinct and mechanistically underexplored form of skeletal disease that is enriched in hormone receptor-positive breast cancers. Preclinical models of bone metastasis from breast cancer predominantly rely on immunodeficient hosts inoculated with osteolytic human breast cancer cell lines, limiting the study of immune-dependent mechanisms of bone remodeling. Here we describe the development and characterization of an immunocompetent, syngeneic osteoblastic bone metastasis model using intratibial injection of PyMT-CK(OB), a luciferase-expressing derivative of the MMTV-PyMT mammary carcinoma cell line, in FVB/N mice.

PyMT-CK(OB) cells produced detectable bioluminescent signal after intratibial injection, enabling longitudinal monitoring of tumor progression. Micro-computed tomography (µCT) revealed significant increases in trabecular bone volume fraction and trabecular number at three and four weeks post-injection, consistent with osteoblastic remodeling. Histological analysis confirmed dense bone lesion formation in tumor-bearing bones. Critically, this osteoblastic phenotype was entirely absent in immunodeficient NOD SCID hosts, despite robust tumor growth, supporting a role for immune competence in tumor-induced bone formation. Loss of bioluminescent signal in immunocompetent mice reflected either immune pressure on reporter gene expression or limited space for cancer cell expansion in the bone, rather than tumor regression or hypoxia, as confirmed by hypoxia imaging and histological endpoint analysis.

In contrast, a second PyMT cell subline, PyMT-CF, maintained sustained bioluminescent signal and produced predominantly osteolytic lesions, providing a complementary syngeneic model of osteolytic disease from the same parental background. In vitro hydrogel coculture experiments and protein array analysis of conditioned media revealed that the PyMT sublines have differing impact on MC3T3 osteoblast mineralization, identifying candidate mediators of divergent bone remodeling phenotypes. R7 mammary carcinoma cells derived from MMTV-RON transgenic mouse mammary tumors did not induce measurable bone remodeling under equivalent experimental conditions. Together, these models provide a validated, immunologically intact framework for studying the mechanistic basis of osteoblastic bone metastasis and evaluating therapeutic interventions targeting the tumor-bone microenvironment.

## Introduction

Skeletal metastasis is among the most debilitating complications of advanced breast cancer, occurring in approximately 70% of patients with metastatic disease and contributing substantially to morbidity through pain, pathological fracture, spinal cord compression, and hypercalcemia [1]. Bone metastases are broadly classified by their net effect on skeletal architecture: osteolytic lesions, driven by tumor-induced osteoclast activation and bone resorption, and osteoblastic lesions that are characterized by excess bone formation stimulated by tumor-derived osteogenic signals [2, 3]. Mixed lesions exhibiting both osteolytic and osteoblastic features are also common, though osteolytic metastases are dominant across breast cancer subtypes [4, 5]. The clinical management of osteoblastic disease remains challenging, and the molecular mechanisms by which tumor cells induce pathological bone formation are incompletely understood.

Osteolytic lesions arise primarily through tumor-induced activation of osteoclasts, the bone-resorbing cells of myeloid lineage, which are recruited and differentiated by receptor activator of NF-κB ligand (RANKL) and macrophage colony-stimulating factor (M-CSF) [6, 7]. During metastatic colonization, tumor cells release factors including parathyroid hormone-related protein (PTHrP) and TGF-β to activate bone forming osteoblasts, leading to RANKL release and activation of osteoclasts, entering a vicious feedback loop in which bone resorption releases matrix-bound growth factors that further fuel tumor proliferation and invasion [8]. Osteoblastic lesions, by contrast, reflect a net shift toward bone formation driven by tumor-derived factors that stimulate osteoblast differentiation through BMP and Wnt signaling pathways and the transcription factor RUNX2, though the tumor-derived signals responsible for initiating this shift remain incompletely characterized [2, 9]. The immune microenvironment is an active participant in regulating bone remodeling. For instance, CD4+ T cells suppress osteoclast formation and restrain resorption [10], myeloid-derived suppressor cells function as direct osteoclast progenitors [11], and neutrophils can directly inhibit osteoblast differentiation to favor net bone loss [12].

Experimental models of breast cancer metastasis to bone have been instrumental in defining the cellular and molecular drivers of skeletal colonization. The most widely used approaches involve intracardiac or intra-arterial injection of human breast cancer cell lines, most commonly MDA-MB-231, into immunodeficient hosts, producing osteolytic lesions detectable by radiographic and histological analysis [3, 13]. Intratibial injection offers a complementary approach that bypasses the metastatic cascade and enables direct, reproducible interrogation of tumor-bone interactions within the marrow niche [3, 14]. However, both approaches are commonly implemented in immunodeficient hosts to accommodate human cell lines, precluding the study of immune contributions to bone remodeling. This is a meaningful limitation because the immune microenvironment, including T cells and myeloid populations, plays an active role in regulating osteoclast and osteoblast activity during metastatic colonization [11, 15, 16]. Syngeneic models in immunocompetent hosts offer a path to studying these interactions, but validated syngeneic models of osteoblastic bone metastasis remain scarce.

The MMTV-PyMT transgenic mouse model drives mammary tumor development under the mouse mammary tumor virus promoter and recapitulates key features of luminal-like hormone receptor-positive human breast cancer, including the propensity for bone metastasis, which is unusual in syngeneic mouse mammary cancer models [17–20]. Cell lines derived from MMTV-PyMT mammary tumors can be injected into syngeneic FVB/N mice, preserving a fully intact immune system. Prior work from our laboratory established that the chemokine ligand CXCL5 and its receptor CXCR2 are sufficient to activate PyMT mammary tumor cell escape from dormancy in bone, using an ex vivo bone colonization system [21]. These findings motivated the development of in vivo intratibial injection models capable of capturing host immune status-dependent bone remodeling.

Here we describe the development of PyMT-CK(OB), a lentiviral luciferase-expressing MMTV-PyMT subline optimized for intratibial injection and bioluminescent monitoring in FVB/N mice. We demonstrate that PyMT-CK(OB) intratibial injection produces reproducible osteoblastic bone lesions as characterized by µCT and histology, and that this phenotype involves an intact immune system. We compare the osteoblastic PyMT-CK(OB) model to a second subline, PyMT-CF, which generates osteolytic lesions, providing complementary syngeneic models from the same parental background. Additional comparisons with R7 mammary carcinoma cells, which are derived from the MMTV-RON model, demonstrate the cell line-induced specificity of the osteoblastic phenotype. We also evaluate intracardiac injection and ex vivo bone co-culture as complementary experimental systems and discuss the practical considerations governing model selection for bone metastasis research.

## Results

### PNA-Luc-EZ BoM cells undergo immune clearance after intratibial injection in immunocompetent FVB/N mice

To model breast cancer metastasis to bone *in vivo*, we first evaluated PNA-Luc-EZ BoM cells, a murine PyMT mammary carcinoma cell line that previously was engineered to stably express GFP and firefly luciferase [22], by intratibial (IT) injection in FVB mice (Figure 1A). Bioluminescence imaging (BLI) of mice detected luciferase signal in tumor-bearing limbs in the first weeks after injection, but signal intensity declined progressively to below background by approximately four weeks (Figure 1B).

**Figure 1.**
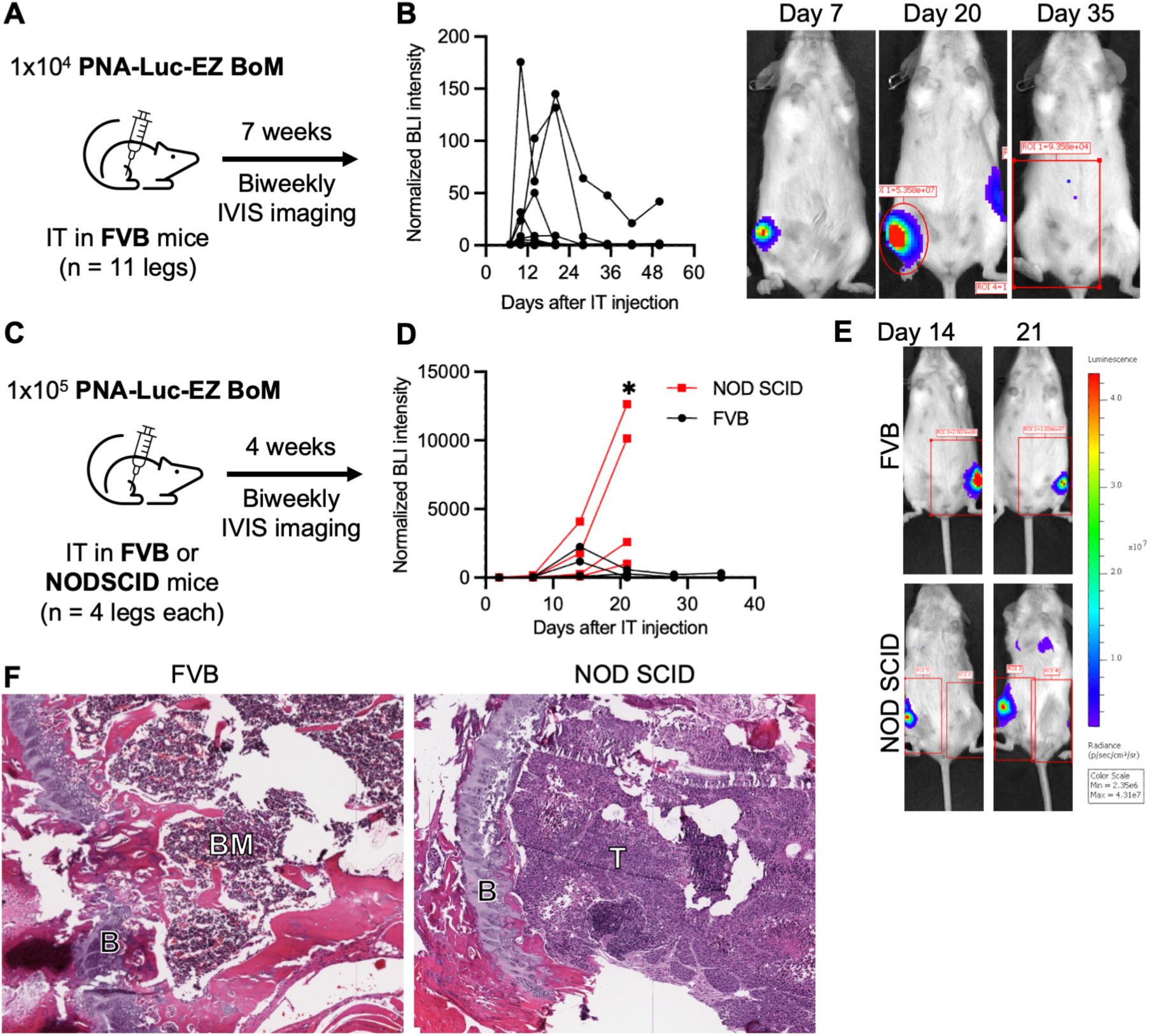
Evaluation of PNA-Luc-EZ BoM IT injection as a bone metastasis model. (A) 1x10^4^ PNA-Luc-EZ BoM cells injected IT into FVB mice (N=11 legs). (B) Quantification and representative IVIS images of lost bioluminescence signal. Data is photon flux in leg ROI normalized to Day 7 BLI. (C) 1x10^5^ PNA-Luc-EZ BoM cells injected IT into FVB or NOD SCID mice (N=4 legs each). (D) Biweekly IVIS imaging to quantify PyMT bioluminescence. Data shown is photon flux in leg ROI normalized to Day 2 BLI. Statistical analysis of FVB vs. NOD SCID at Day 21 by Mann-Whitney test, P=0.0286. (E) Representative IVIS images of PNA-Luc-EZ BoM IT FVB and NOD SCID mice. (F) H&E staining of IT-injected bone marrow in the proximal tibia of FVB and NOD SCID mice. B = bone, growth plate in proximal tibia; BM = bone marrow; T = tumor cells. Significance levels: * p < 0.05.

To determine whether this loss reflected an immune response to the cancer cells expressing the luciferase and GFP reporter genes, rather than turning off the reporter genes to allow tumor growth that could not be detected by IVIS, we compared tumor progression in FVB/N to tumor progression in NOD SCID immunodeficient mice after 1x10^5^ PNA-Luc-EZ BoM cell IT injection (Figure 1C). In NOD SCID mice, BLI signal increased continuously throughout the study period, and tumors progressed to lung metastasis, necessitating early euthanasia in a subset of animals (Figure 1D, E). In contrast, FVB/N mice again exhibited transient signal followed by complete loss.

Endpoint histological analysis of hematoxylin and eosin (H&E)-stained bones confirmed extensive metastatic colonization in NOD SCID tibiae, while FVB/N bones showed no residual tumor cells and intact bone marrow architecture, suggesting that an immune response cleared the cancer cells, rather than turning off the luciferase and GFP reporter genes and leaving cancer cells (Figure 1F). These findings indicate that PNA-Luc-EZ BoM cells are cleared from the bone marrow of immunocompetent FVB/N mice, possibly due to immune rejection of the GFP or luciferase transgenes.

### Lentiviral transduction of PyMT-CL cells produces PyMT-CK(OB), a luciferase-expressing subline with a detectable in vivo signal

Reporter gene immunogenicity in immunocompetent hosts has been documented for GFP and high-level luciferase expression [23, 24]. We therefore evaluated a GFP-negative, luciferase-positive PyMT cell line (PyMT-CL) that was obtained from Dr. Conor Lynch (Moffitt Cancer Center) [20]. The cell culture BLI confirmed that PyMT-CL cell luminescence was substantially dimmer than in PNA-Luc-EZ BoM cells (Figure 2A). Intratibial injection of 1x10^5^ PyMT-CL cells in FVB/N mice precluding temporal longitudinal monitoring of tumor growth (Figure 2B).

**Figure 2.**
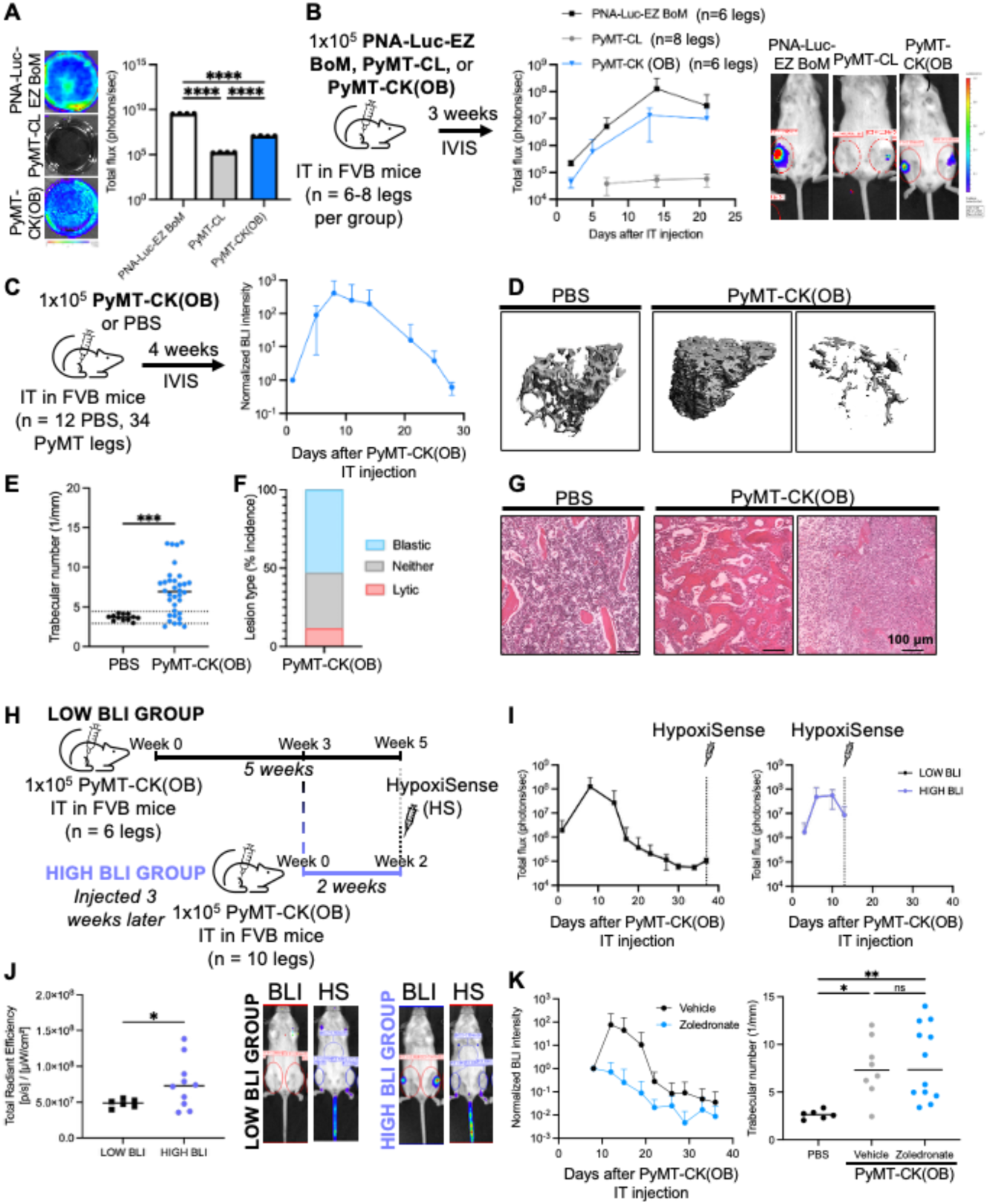
Characterization of luciferase+ PyMT (PyMT-CK(OB)) cell bioluminescence and bone remodeling after IT injection. (A) Luminescence intensity of PNA-Luc-EZ BoM, PyMT-CL, and PyMT-CK(OB) cells in culture. Statistical analysis by ANOVA, p<0.0001. (B) 1x10^5^ PNA-Luc-EZ BoM, PyMT-CL, or PyMT-CK(OB) cells injected IT into FVB mice (N=6-8 legs per group). PyMT-CL tumor growth was not detected by BLI, while PyMT-CK(OB) tumors have an increased BLI signal in vivo. IVIS was used to quantify tumor bioluminescence for leg ROI. Data shown is mean ± SD for total photon flux and representative IVIS images of IT tumors on day 21. (C) 1x10^5^ PyMT-CK(OB) cells (N=34 legs) or PBS (N=12 legs) were injected IT in FVB mice and imaged biweekly for four weeks. Data shown is mean ± SD for photon flux in leg ROI normalized to day 1 signal. (D) Representative 3D reconstructions and (E) μCT quantification of trabecular number in PBS and PyMT-CK(OB) tumor-bearing female FVB mouse bones. Solid lines indicate mean, dashed lines indicate 2 SD from the mean PBS Tb. N, used to categorize osteoblastic and osteolytic lesions. Statistical analysis by Mann-Whitney, P=0.0007. (F) Incidence of osteoblastic and osteolytic remodeling in PyMT-CK(OB) IT bone lesions. (G) H&E staining of medullary cavity in the proximal tibia of healthy (PBS IT) and PyMT-CK(OB) tumor-bearing mice. PyMT-CK(OB) IT injection produces predominantly osteoblastic bone lesions. (H) HypoxiSense probe imaging of FVB mice after two weeks (n=10 legs) or five weeks (n=6 legs) following 1x10^5^ PyMT-CK(OB) cells IT injection. (I) BLI intensity showing timing of luminescence and imaging. Data shown is mean ± SD for total photon flux in leg ROI. (J) Fluorescence intensity (total radiant efficiency) in leg ROI of low and high BLI signal groups and representative IVIS images of PyMT-CK(OB) luminescence and HypoxiSense fluorescence in low BLI (5 weeks post-IT) and high BLI (2 weeks post-IT) groups. Lines indicate the mean. Statistical analysis by Welch’s t-test (P=0.0352). (K) 1x10^5^ PyMT-CK(OB) cells were injected IT in FVB mice and mice were treated with zoledronate or vehicle control 3x/week s.c. Data shown is mean ± SD for photon flux in leg ROI normalized to day 8 treatment start signal and μCT quantification of trabecular number in PBS and vehicle- or zoledronate-treated PyMT-CK(OB) tumor-bearing female FVB mouse bones. Significance levels: * p < 0.05, **p < 0.01, ***p < 0.001, ****p < 0.0001, ns: not significant (p ≥ 0.05).

To increase bioluminescent signal while retaining the PyMT-CL genetic background, we transduced the PyMT-CL cells with a lentiviral luciferase expression vector (Lenti-luciferase-P2A-Neo) followed by neomycin selection to generate PyMT-CK(OB) cells (named OB for <u>o</u>steo<u>b</u>lastic activity described below). BLI of PyMT-CK(OB) cells in culture confirmed a significant increase in luminescence relative to the parental PyMT-CL line (Figure 2A).

Following intratibial (IT) injection in FVB/N mice, PyMT-CK(OB) cells produced in vivo BLI signal of intermediate intensity between that of PyMT-CL and PNA-Luc-EZ BoM, with tumors detectable within days of injection (Figure 2B).

### PyMT-CK(OB) intratibial injection induces osteoblastic bone remodeling in immunocompetent FVB/N mice

To characterize the bone remodeling response to PyMT-CK(OB) during colonization, FVB/N mice received IT injections of 1x10^5^ PyMT-CK(OB) cells or PBS, and bones were collected at four weeks (Figure 2C). BLI signal from PyMT-CK(OB) bones peaked at approximately two weeks after IT injection before progressively declining.

To formally classify bone remodeling phenotypes as osteoblastic or osteolytic, μCT analysis was used to quantify trabecular number (Tb.N) values from four-week PyMT-CK(OB) IT bones compared to PBS controls (Figure 2D-E). The mean Tb.N was 3.69 ± 0.39 in PBS and 6.94 ± 3.06 in PyMT-CK(OB) bones (Figure 2E, N=12 PBS and 34 PyMT-CK(OB) bones, P=0.0007, Mann-Whitney). Lesions were defined as osteoblastic or osteolytic based on Tb.N values exceeding or below two standard deviations from the PBS IT control mean, respectively. From this analysis, the majority of PyMT-CK(OB) IT lesions were classified as osteoblastic (mean Tb.N elevated versus PBS), with a subset exhibiting osteolytic lesions (Figure 2F).

H&E histological analysis of four-week old bones confirmed the μCT classifications (Figure 2G). Thus, the decline in BLI intensity at later tumor timepoints is likely attributable in part to limited space in and attenuation of the luminescent signal through the increasingly dense bone tissue [25], rather than immunological tumor clearance. While bones with PBS injection preserved the normal bone marrow morphology, osteoblastic PyMT-CK(OB) bones showed dense trabecular lesions replacing normal marrow, and osteolytic bones showed cortical and trabecular degradation. These findings establish PyMT-CK(OB) IT injection in FVB/N mice as a reproducible syngeneic model of predominantly osteoblastic breast cancer bone metastasis.

### Hypoxia does not account for the progressive bioluminescence signal loss seen in PyMT-CK(OB) intratibial tumors

We next evaluated hypoxia, immune clearance, and space constraints as reasons why the luciferase signal declines temporally. Firefly luciferase activity requires D-luciferin and molecular oxygen as the substrate to generate oxyluciferin and yellow-green light [26], raising the possibility that hypoxia accumulating in the bone tumor microenvironment could limit BLI signal at later timepoints. To evaluate this, we tested for hypoxic environments in tumors that had high or low BLI using a fluorescent probe for hypoxia. We administered the IVISense Hypoxia CA IV 680 fluorescent probe (HypoxiSense), which detected carbonic anhydrase IX (CAIX) as a surrogate of regional hypoxia, via tail vein injection in mice bearing PyMT-CK(OB) IT tumors at either peak BLI (high BLI signal, approximately two weeks post-injection, N=10 legs) or declining BLI (low signal, approximately five weeks post-injection, N=6 legs) timepoints (Figure 2H-J).

Fluorescence imaging revealed minimal hypoxia signal accumulation in the tibial tumor region (Figure 2J). Quantification showed significantly higher hypoxia signal in the high-BLI group relative to the low-BLI mice (Figure 2J, P=0.0352; Welch’s t-test), the opposite of what would be expected if oxygen limitation were driving BLI signal loss.

These data indicate that progressive BLI decline in PyMT-CK(OB) IT tumors is not attributable to tumor hypoxia.

### PyMT-CK(OB)-induced osteoblastic remodeling is not prevented by zoledronate treatment

We also tested whether osteoblastic PyMT-CK(OB) bone tumors would respond to the clinically used bone resorption inhibitor zoledronate after intratibial injection (Figure 2K). Zoledronate treatment significantly slowed the progression of PyMT-CK(OB) tumors by BLI (Fisher’s combined P = 0.0156). However, μCT analysis showed that dense osteoblastic lesions characterized by a significantly increased Tb.N compared to PBS controls were not prevented by zoledronate treatment (vehicle versus zoledronate P = 0.9242). These data indicate that while zoledronate may act through a direct tumor-targeted mechanism to slow tumor growth [27], it does not prevent PyMT-CK(OB) cell-induced bone formation after intratibial injection.

### Concurrent osteoblastic bone remodeling and BLI signal loss in FVB/N mice, but not in NOD SCID mice, support an immune-dependent mechanism of PyMT-CK(OB)-induced bone remodeling that is distinct from complete tumor cell clearance

A possible explanation for the loss of BLI intensity in PyMT-CK(OB) IT bones is that tumor cells are cleared from bone due to an immune response to luciferase, as seen in the PNA-Luc-EZ BoM cells with GFP [23, 24], or that the osteoblastic remodeling in FVB/N mice limits cancer cell expansion in the bone. To determine whether an immune response drives both BLI signal loss and osteoblastic bone remodeling, we compared FVB/N and NOD SCID mice receiving IT injection of PBS or PyMT-CK(OB) cells and collected bones at two, three, and four weeks (Figure 3A).

**Figure 3.**
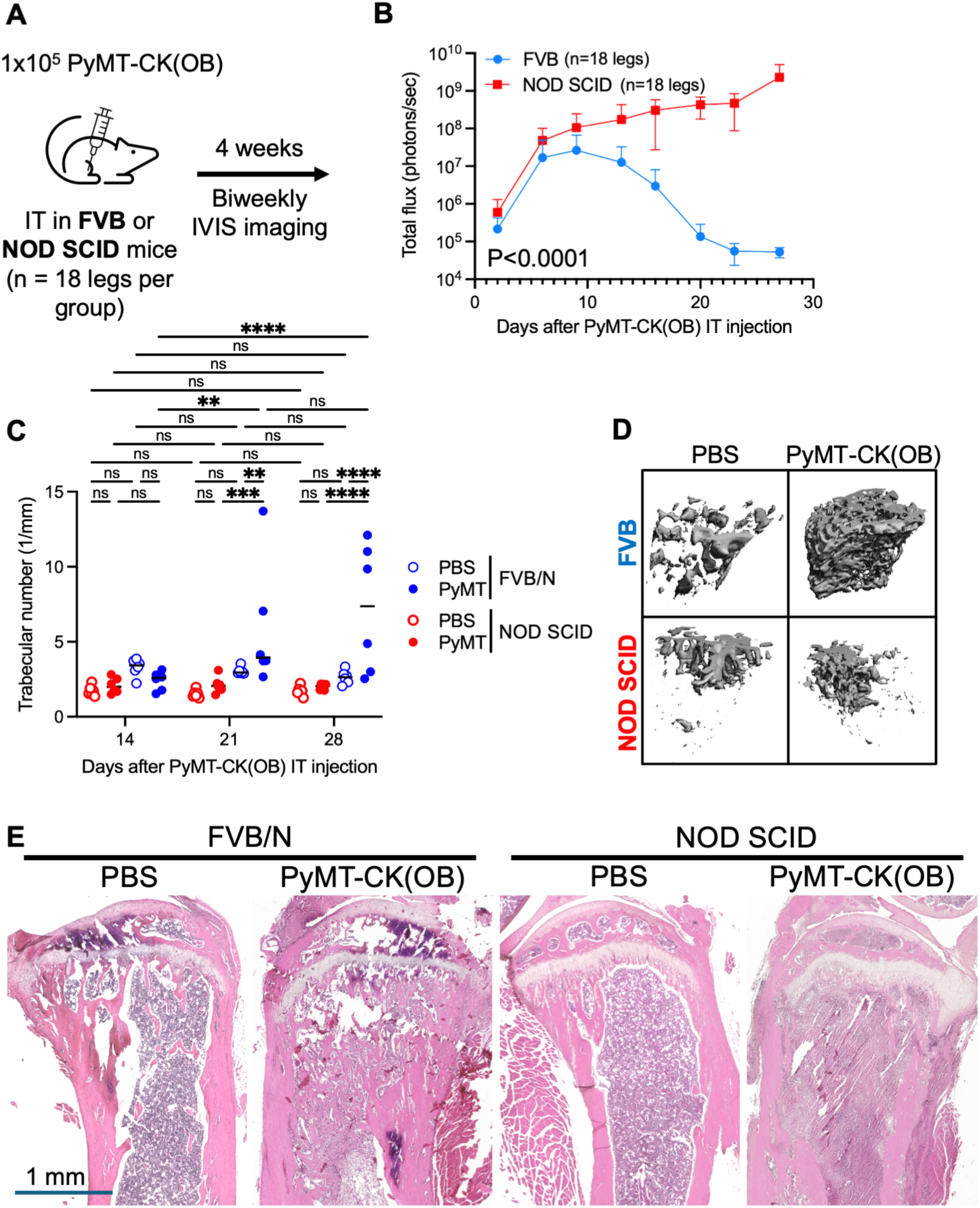
Host immune competence is associated with osteoblastic remodeling and reduced cancer cell BLI after PyMT-CK(OB) intratibial injection. (A) 1x10^5^ PyMT-CK(OB) cells IT in FVB and NOD SCID mice (N=36 legs each), collected two, three, or four weeks after injection (N=12 legs/timepoint/strain) for μCT. (B) BLI intensity in FVB mice drops after ∼two weeks, while PyMT-CK(OB) signal in NOD SCID increases for the study duration. Data shown is mean ± SD for total photon flux. (C) Trabecular number quantification and (D) 3D reconstruction in the proximal tibia of FVB and NOD SCID mice. PyMT-CK(OB) IT injection induced significant osteoblastic bone remodeling in FVB mice and no changes in NOD SCID. Statistical analysis by ANOVA. Significance levels: **p < 0.01, ***p < 0.001, ****p < 0.0001, ns: not significant (p ≥ 0.05). (E) Histology by H&E staining shows normal bone marrow morphology in PBS bones and osteoblastic remodeling in FVB/N, but not NOD SCID, tibiae after PyMT-CK(OB) IT injection.

While NOD SCID mice do not produce T or B cells, they also have defective complement and NK cell responses and reduced innate immunity through impaired dendritic cell maturation and defective macrophage populations [28–30]. BLI monitoring confirmed the strain-dependent divergence in signal kinetics: FVB/N mice showed signal decline after approximately two weeks, while NOD SCID mice exhibited continuous signal over the full four-week period (Figure 3B). This suggests that the BLI decline could be due either to an immune response that kills the cancer cells or limits the luciferase accumulation or to a physical restriction of the infiltration of the luciferase-expressing cancer cells to the bone. To further distinguish these possibilities, we evaluated the bones by μCT analysis and by histology.

μCT analysis revealed a striking contrast in bone architecture between the two host strains (Figure 3C-D). In FVB/N mice, PyMT-CK(OB) IT injection produced robust osteoblastic remodeling, with significantly elevated Tb.N in tumor-bearing bones compared to PBS at three and four weeks (week 4 Tb.N = 2.63 ± 0.45/mm in PBS vs. 7.23 ± 4.25/mm in PyMT-CK(OB)). In contrast, NOD SCID mice showed no significant difference in PyMT-CK(OB) trabecular number compared to PBS controls at any timepoint (week 4 Tb.N = 2.00 ± 0.21/mm in PBS vs. 1.73 ± 0.32/mm in PyMT-CK(OB)). This suggests that the osteoblastic remodeling requires immune competence, consistent with published studies demonstrating that T cells and myeloid populations regulate tumor-induced bone remodeling in syngeneic models [15, 16].

Histological analysis with H&E staining of PyMT-CK(OB) IT FVB bones confirmed that cancer cells were present and displayed formation of dense bone lesions compared to healthy bone marrow, even though the BLI signal had declined (Figure 3E). H&Es of NOD SCID bones show extensive colonization by PyMT-CK(OB) cells with no apparent osteoblastic remodeling.

Together, these data demonstrate that accumulation of luciferase-expressing cancer cells alone is not sufficient to drive the bone remodeling and suggests that functional immune activity is involved in PyMT-CK(OB)-induced osteoblastic bone remodeling. In addition, the BLI signal loss in PyMT-CK(OB) tumors occurs only with an intact immune response, suggesting either that a physical restriction from the bone remodeling prevents the cancer cell infiltration or that an immune response eliminates the reporter-expressing tumor cells.

### PyMT-CF cells maintain BLI signal and induce osteolytic bone remodeling after IT injection

The progressive BLI signal loss in PyMT-CK(OB) cells complicates longitudinal tumor monitoring in FVB/N mice. We generated a second luciferase-expressing PyMT-CL subline, PyMT-CF, by independent lentiviral transduction and neomycin selection. BLI of PyMT-CF cultured cells confirmed that the luminescence of PyMT-CF cells was comparable to that of PyMT-CK(OB) cells (Figure 4A).

**Figure 4.**
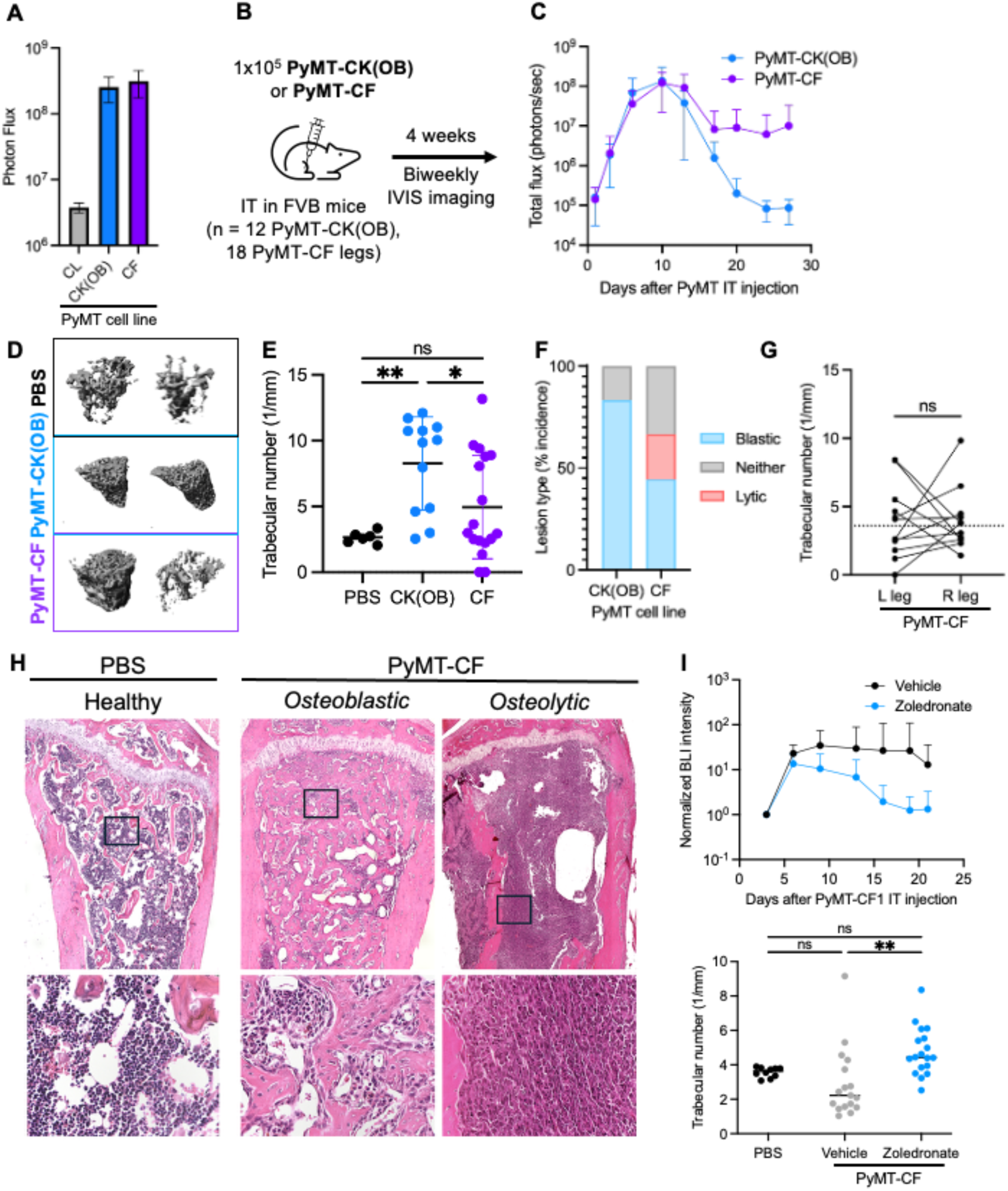
Characterization of luciferase-expressing PyMT-CF cell lines that have osteolytic activity. (A) Luminescence intensity mean ± SD of PyMT-CL, -CK(OB), and -CF cells in culture. (B) 1x10^5^ PyMT-CK(OB) cells (n=12 legs) or PyMT-CF cells (n=18 legs) were injected IT into FVB mice. (C) Quantification of tumor growth by BLI of PyMT-CK(OB) and PyMT-CF bone tumors for four weeks. Data shown is mean ± SD for total photon flux in leg ROI. (D) Representative μCT 3D reconstructions and (E) quantification of trabecular number in bones four weeks after IT injection of PBS, PyMT-CK(OB), or PyMT-CF. (F) Incidence of osteoblastic and osteolytic lesions compared to PBS bones in PyMT-CK(OB) and PyMT-CF bones. Statistical analysis by Fisher’s exact test, P=0.0504. (G) Quantification of trabecular number in paired L and R leg bones from PyMT-CF IT mice, showing osteoblastic and osteolytic tumors in the same mouse. Dotted line indicates the mean trabecular number in PBS IT mice. Statistical analysis by Wilcoxon test, P=0.8501. (H) H&E staining of PyMT-CF tumor-bearing bones displaying normal bone marrow compared to abnormal bone formation or severe bone degradation. (I) 1x10^5^ PyMT-CF cells were injected IT in FVB mice and mice were treated with zoledronate or vehicle control 3x/week s.c. Data shown is mean ± SD for photon flux in leg ROI normalized to day 3 treatment start signal and μCT quantification of trabecular number in PBS and vehicle- or zoledronate-treated PyMT-CF tumor-bearing female FVB mouse bones. Significance levels: ns: not significant (p ≥ 0.05), * p < 0.05, ** p < 0.01.

We next compared the in vivo behavior of these cell lines after intratibial injection into FVB/N mice. Interestingly, all PyMT-CF cells maintained substantially higher BLI signal over time compared to PyMT-CK(OB), suggesting this cancer cell subline experiences either reduced immune pressure or fewer physical constraints within the bone microenvironment (Figure 4B-C). We evaluated the bones by μCT analysis and by histology.

MicroCT analysis of the bones four weeks after IT injection of 1x10^5^ PyMT-CF cells revealed a markedly different bone remodeling phenotype compared to PyMT-CK(OB) cells, wherein PyMT-CF bones exhibited mixed and frequently osteolytic lesions, in contrast to the predominantly osteoblastic phenotype of PyMT-CK(OB) (Figure 4D–F). In fact, several bones with PyMT-CF tumors were too severely degraded to permit standardized μCT contouring (depicted as Tb.N = 0), indicating advanced osteolysis.

To characterize if the heterogeneity of bone remodeling responses is inherent to the cell lines, we analyzed tumors produced from paired tibiae from within mice receiving bilateral intratibial injections with 1x10^5^ PyMT-CF cells. Both osteoblastic and osteolytic lesions were evident across the cohort, and in some mice, left and right tibiae exhibited opposite remodeling phenotypes (Figure 4G). This within-mouse heterogeneity indicates that the cells have the potential to become either osteoblastic or osteolytic to govern the net direction of tumor-induced remodeling.

H&E analysis of the bones that could be analyzed confirmed extensive destruction of trabecular and cortical architecture in osteolytic PyMT-CF IT bones (Figure 4H).

### PyMT-CF tumors are responsive to zoledronate treatment after intratibial injection

We also tested the response of PyMT-CF tumors to treatment with the bone resorption inhibitor zoledronate after intratibial injection (Figure 4I). Zoledronate significantly reduced the growth of PyMT-CF tumors by BLI (Fisher’s combined P<0.0001). MicroCT analysis demonstrated that osteolytic remodeling induced by PyMT-CF tumor growth was significantly inhibited by zoledronate treatment, since vehicle-treated bones had a significantly lower trabecular number compared to treated bones (P=0.0028). These data support that PyMT-CF tumors drive an osteoclast-mediated bone remodeling that is distinct from that of PyMT-CK(OB) cells.

### Distinct bone remodeling programs from osteoblasts cocultured with the PyMT-CK(OB) and PyMT-CF sublines

Even though the PyMT-CK(OB) and PyMT-CF cells are derived from the same PyMT-CL cell line, these sublines revealed different potentials to develop into osteoblastic and osteolytic bone metastasis. Interested in the predominantly osteoblastic phenotype of the PyMT-CK(OB) cells and the comparatively more osteolytic yet more heterogeneous PyMT-CF cells, we analyzed the functional differences of these cells on bone remodeling using coculture experiments. We utilized the previously developed in vitro three-dimensional model coculturing PyMT cancer cells and MC3T3 murine osteoblasts embedded in separate gelatin hydrogels crosslinked with transglutaminase [31]. This model enables paracrine signaling between the osteoblasts and cancer cells while maintaining a physiologically relevant microenvironment. MC3 osteoblasts were cocultured with PyMT cancer cells (MC3+PyMT-CK(OB), MC3+PyMT-CF) for 28 days in osteogenic media (Figure 5A).

**Figure 5.**
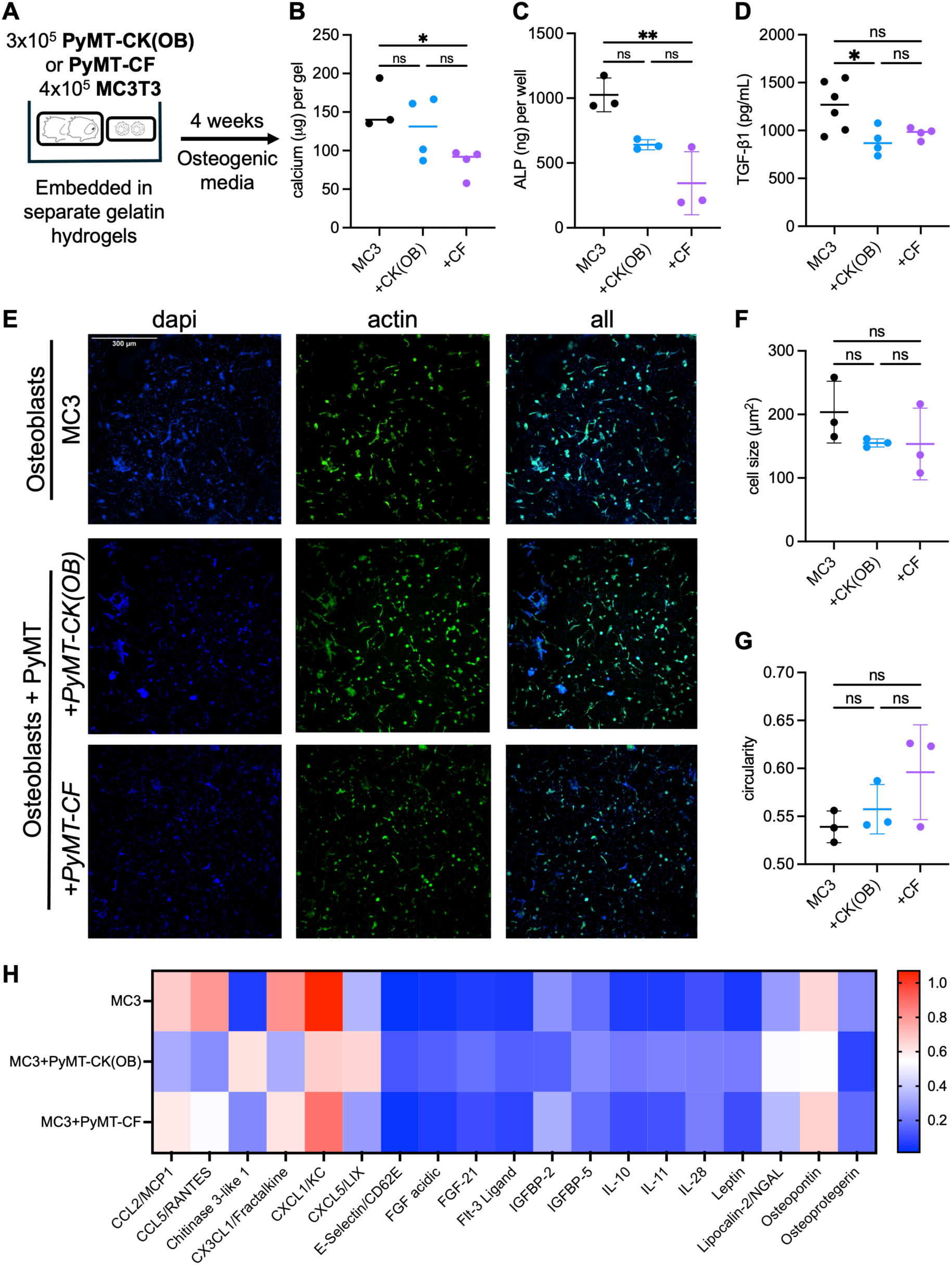
**Evaluation of bone remodeling in in vitro coculture with PyMT cells**. (A) Experimental design indicating 3 × 10^5^ PyMT cells were embedded into hydrogels and cultured together with 4 × 10^5^ MC3T3 cells in a separate hydrogel for 4 weeks with osteogenic supplementation. Quantification of (B) bone mineral deposition by calcium assay and (C) osteoblast maturation by ALP assay. (D) Quantification of TGF-β1 by ELISA of culture media. (E) Representative immunofluorescent nuclear and cytoskeletal imaging of MC3T3 cells in hydrogels in MC3, MC3+PyMT-CK(OB), and MC3+PyMT-CF groups and quantification of (F) cell size and (G) cell circularity. Lines indicate mean ± SD. (H) Protein array analysis reveals distinct secretion in PyMT-CK(OB) cocultures compared to MC3 monoculture and PyMT-CF coculture. Statistical analysis by ordinary one-way ANOVA. Significance levels: ns: not significant (p ≥ 0.05), * p < 0.05, ** p < 0.01.

During bone formation, osteoblasts first synthesize, secrete, and lay down extracellular matrix (ECM; osteoid) and then load the osteoid with minerals (Ca^2+^ and P_i_ ions that precipitate and form hydroxyapatite crystals; mineralization). We measured by calcium assay the mineralization of MC3 cells grown in coculture with either PyMT-CK(OB) or PyMT-CF cells (Figure 5B). When cocultured with PyMT-CK(OB) cells, MC3 mineralization was unaffected. In contrast, being cocultured with PyMT-CF cells inhibited MC3 mineralization. Similarly, alkaline phosphatase (ALP) secretion was not inhibited in PyMT-CK(OB) cocultures but was inhibited in PyMT-CF cocultures (Figure 5C). These results highlight functional mineralization differences between the PyMT-CK(OB) and PyMT-CF cells, wherein PyMT-CK(OB) cells did not directly stimulate bone mineralization, while PyMT-CF cells suppress osteoblast mineralization and maturation, consistent with an osteolytic phenotype.

The lack of increased osteoblastogenesis by the PyMT-CK(OB) coculture suggests either that osteoblastic lesion formation may require other stromal or marrow cell involvement or that increased osteogenic supplementation in the in vitro models may be required to support the increased osteogenesis. Indeed, the osteogenic factor TGF-β1 levels were lower in PyMT-CK(OB) coculture compared to MC3 groups (Figure 5D).

Osteoblasts in coculture with either PyMT-CK(OB) or PyMT-CF cancer cells were not significantly different in cell size or circularity compared to osteoblasts alone (Figure 5E-G), indicating that coculture with either PyMT subline did not halt osteoblast differentiation in three-dimensional culture.

### Protein array analysis of conditioned media highlights distinct secretions of pro-inflammatory and osteogenic factors from cocultures of the PyMT-CK(OB) and PyMT-CF sublines

To investigate the osteogenic factors that could contribute to the osteogenic reprogramming from the PyMT sublines, we analyzed the conditioned media for osteogenic factors, including 111 soluble mouse cytokines and chemokines, by protein array (Proteome Profiler Mouse XL Cytokine Array, R&D Systems) (Figure 5H). Protein analysis of coculture media revealed that PyMT-CK(OB) cells had elevated levels of some pro-inflammatory cytokines, including chitinase 3-like 1, E-selectin, IL-10, IL-11, and CXCL5/LIX. These factors are consistent with the involvement of immune cell populations in the osteoblastic remodeling derived from PyMT-CK(OB) cells. The absence of immune cells in this in vitro model likely contributes to the lack of increased osteoblastic behavior of the MC3T3 cells.

PyMT-CK(OB) coculture also exhibited reduced levels of several proteins including CCL2/MCP1 and osteoprotegerin, indicating reduced osteoclastogenic signaling by osteoblasts [32, 33], which could further exacerbate blastic lesion formation.

### Intracardiac injection delivery of PyMT-CK(OB) cells does not produce osteoblastic bone remodeling

To determine whether the osteoblastic remodeling of PyMT-CK(OB) cells depends on a tumor cell intrinsic property or on the route for local delivery to the bone marrow niche during systemic metastasis, we compared bone remodeling in mice receiving intratibial versus intracardiac injection of PyMT-CK(OB) cells (Figure 6A).

**Figure 6.**
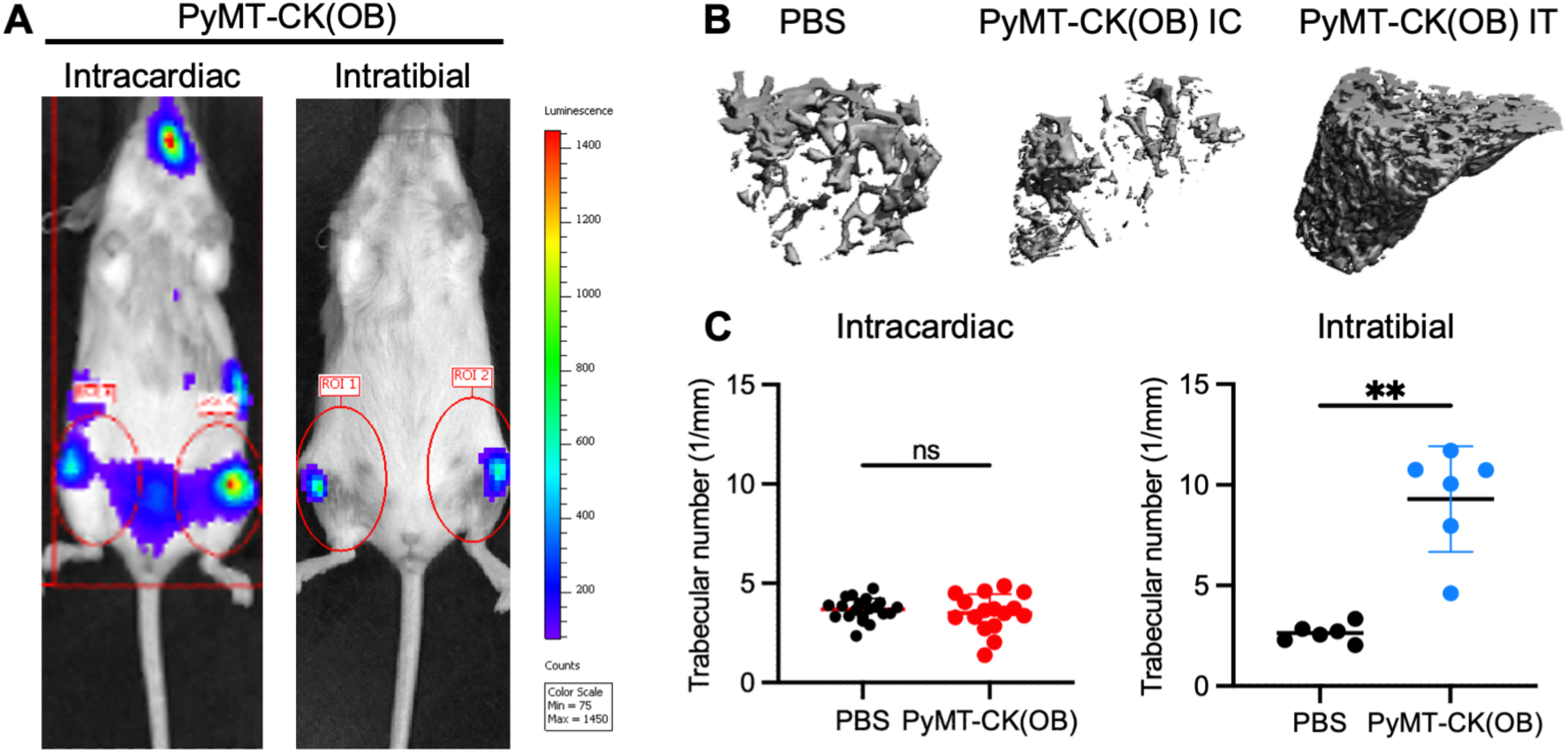
Bone remodeling after PyMT-CK(OB) tumor growth depends on injection site. (A) Bioluminescent imaging of mice after intracardiac (IC) or intratibial (IT) injection of 1x10^5^ PyMT-CK(OB) cells. (B) Representative 3D reconstructions and (C) trabecular number quantification of healthy or tumor-bearing bones from μCT analysis. Lines indicate mean ± SD. Statistical test of PBS vs. PyMT-CK(OB) by Mann-Whitney. Significance levels: **p < 0.01, ns: not significant (p ≥ 0.05).

BLI following intracardiac injection confirmed tumor cell distribution to multiple organs, including lung, kidney, ovary, and bone. µCT analysis of tibiae from intracardiac-injected mice were not significantly different in trabecular bone architecture relative to controls, in contrast to the osteoblastic remodeling observed with intratibial injection (Figure 6B-C). These findings could suggest that a direct local interaction between PyMT-CK(OB) cells in clusters or between PyMT-CK(OB) and the bone microenvironment may be required for the osteoblastic bone remodeling or could implicate the wound-healing response to the physical wound of IT injection as a contributing factor to bone formation.

### R7 mammary carcinoma cells colonize bone but do not induce bone remodeling

R7 cells are a murine mammary carcinoma line derived from MMTV-RON transgenic tumors that express estrogen receptor and respond to tamoxifen in culture [34, 35]. To evaluate their bone metastasis potential, FVB/N mice received IT injection of 1x10^5^ R7 cells without a luciferase reporter, and bones were collected two, three, and four weeks post-injection (Figure 7A). µCT analysis showed no significant difference in trabecular number between R7-injected and PBS control bones at any timepoint by two-way ANOVA with Fisher’s LSD (week 2 P=0.074, week 3 P=0.468, week 4 P=0.299; Figure 7B). PBS control Tb.N declined significantly between weeks three and four (P=0.049), consistent with age-related trabecular thinning over the study period rather than tumor-driven remodeling. No osteolytic lesions were identified by µCT classification criteria at any timepoint, and no cortical degradation or severe trabecular loss was observed.

**Figure 7.**
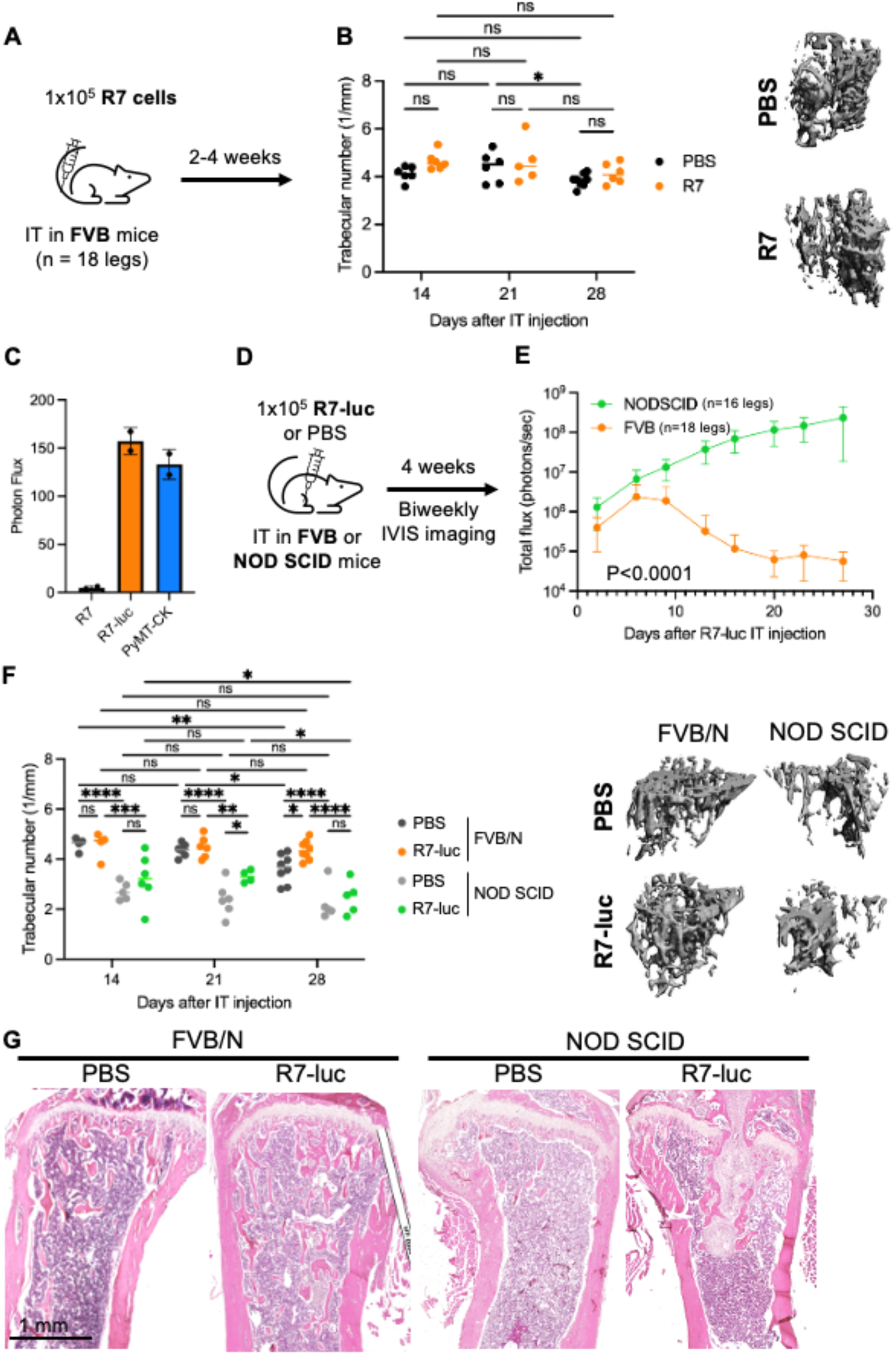
Characterization of R7 cell bone colonization and trabecular remodeling after IT injection. (A) 1x10^5^ R7 cells or PBS were injected IT into FVB mice (n=18 legs each), and bones were assessed by μCT at days 14, 21, and 28. (B) μCT quantification of trabecular number and representative 3D reconstructions of PBS and R7 tumor-bearing female FVB mouse bones at day 28. (C) Luminescence intensity of R7, R7-luc, and PyMT-CK(OB) cell lines in culture. (D) 1x10^5^ R7-luc cells or PBS were injected IT into immunocompetent FVB or immunodeficient NOD SCID mice and imaged biweekly by IVIS for four weeks. (E) Quantification of total bioluminescence flux (photons/sec) over time in FVB (n=18 legs) and NOD SCID (n=16 legs) mice following R7-luc IT injection. Data shown is mean ± SD for total photon flux. R7-luc tumor bioluminescence was significantly higher in NOD SCID mice compared to FVB mice (P<0.0001, mixed-effects analysis). (F) μCT quantification of trabecular number and representative 3D reconstructions of PBS and R7-luc tumor-bearing FVB/N and NOD SCID mouse bones at days 14, 21, and 28. R7-luc IT injection did not produce consistent trabecular bone remodeling in either FVB/N or NOD SCID hosts. Statistical analysis by two-way ANOVA with Fisher’s LSD. (G) Representative H&E staining of PBS control and R7-luc tumor bearing bones from FVB and NOD SCID mice. Significance levels: *p < 0.05, **p < 0.01, ***p < 0.001, ****p < 0.0001, ns: not significant (p ≥ 0.05).

To enable longitudinal monitoring of tumor colonization and to evaluate R7 behavior across host strains, we generated luciferase-expressing R7-luc cells by lentiviral transduction (Figure 7C). The in vitro luminescence of these cells was comparable to PyMT-CK(OB) cells after neomycin selection. FVB/N and NOD SCID mice received IT injection of 1x10^5^ R7-luc cells and were monitored by biweekly BLI for four weeks (Figure 7D). In NOD SCID hosts, BLI signal increased progressively throughout the study period, confirming successful engraftment and sustained tumor growth in bone. In FVB/N hosts, BLI signal declined after an initial peak, consistent with host-mediated reporter attenuation observed across cell lines in this study (Figure 7E).

MicroCT analysis of R7-luc IT bones showed no significant difference in trabecular number between PBS and R7-luc tumor-bearing bones within either host strain at any timepoint, with the exception of a modest increase in R7-luc bones relative to PBS at week four in FVB/N mice (P=0.011) and at week three in NOD SCID mice (P=0.014), both in the osteoblastic, rather than osteolytic, direction (Figure 7F). FVB/N PBS control Tb.N declined significantly between weeks two and four (P=0.006) and weeks three and four (P=0.018), indicating that the week four FVB/N result reflects progressive PBS trabecular thinning, rather than tumor-driven bone formation. NOD SCID R7-luc Tb.N declined modestly but significantly across the study period (week two vs four P=0.030, week three vs four P=0.028), while NOD SCID PBS controls showed no significant change across timepoints. This progressive decline in tumor-bearing NOD SCID bones, in the context of confirmed tumor growth by BLI, may reflect modest osteolytic activity, though within-timepoint PBS versus R7-luc comparisons remained non-significant throughout. Notably, FVB/N PBS controls had significantly higher baseline trabecular number than NOD SCID PBS controls at all three timepoints in this cohort (all p<0.0001), consistent with a directional trend toward lower trabecular bone volume in NOD SCID mice observed in the PyMT-CK(OB) FVB/N versus NOD SCID cohorts. Reduced trabecular bone mass in immunodeficient mice relative to immunocompetent controls has been reported and attributed to altered immune-bone crosstalk in the absence of functional lymphoid populations [36].

Histological analysis of PBS and R7-luc IT-injected bone confirmed that tumor cells were present in FVB/N bone despite loss of BLI signal and lack of detectable bone remodeling (Figure 7G), consistent with host-mediated silencing of reporter gene expression in vivo. Accordingly, increased tumor burden was observed in NOD SCID bones after R7-luc IT injection though with no apparent effect on cortical or trabecular bone structure.

Together, these data demonstrate that under these experimental conditions, bone colonization is not sufficient for tumor-induced bone remodeling and that R7 cells lack the osteoinductive or osteoclastogenic signals necessary to alter trabecular architecture after intratibial delivery, regardless of immune status.

## Discussion

This study created and characterized the intratibial injection of PyMT-CK(OB) cells as a unique and reproducible syngeneic model of osteoblastic breast cancer bone metastasis and bone remodeling, while PyMT-CF is a complementary PyMT model with predominantly osteolytic/mixed remodeling. Osteoblastic lesions, which are associated clinically with hormone receptor-positive disease and represent a mechanistically distinct form of skeletal involvement, currently are substantially underrepresented in preclinical bone metastasis research relative to their osteolytic counterparts. The availability of an immunologically intact syngeneic model that reproducibly generates osteoblastic lesions provides a new experimental platform for dissecting the cellular and molecular interactions between tumor cells, bone-remodeling cells, and the immune microenvironment.

A biologically significant finding of this work is the dependence on host immune competence in PyMT-CK(OB)-induced osteoblastic bone remodeling based on the comparison in FVB/N and NOD SCID hosts. Despite robust increases in tumor growth in immunodeficient NOD SCID bones, these bones produced no detectable change in trabecular architecture. In contrast, immune-intact FVB/N bones bearing comparable tumor burden to in NOD SCID bones developed substantial bone formation. This immune dependence likely reflects the contributions of T cells and other adaptive immune populations as well as the innate immune cell populations –which are defective in these mice –to osteoblast activation during metastatic colonization [15, 16]. These findings also have a direct practical implication for model selection, given that the widespread use of immunodeficient hosts in bone metastasis research, necessitated by the use of human cell lines, precludes the study of these immune-dependent mechanisms.

A major practical challenge encountered in the development of this model was the progressive loss of bioluminescent signal in immunocompetent FVB/N mice, despite confirmed tumor persistence by histology. This phenomenon was observed across multiple PyMT-derived sublines and was absent in NOD SCID hosts. One interpretation of the loss of BLI is an immune-mediated mechanism directed against luciferase or other reporter gene products, rather than true tumor regression or hypoxia-induced enzyme inactivation. Hypoxia imaging with the HypoxiSense probe explicitly ruled out oxygen limitation as a contributor to BLI signal loss. These findings are consistent with published reports of reporter gene immunogenicity in immunocompetent mouse models [23, 24, 37, 38] and highlight a critical limitation of BLI as a sole endpoint for tumor burden assessment in immune-intact bone metastasis models. In this context, endpoint µCT and histological analysis are essential complements to BLI and should be treated as primary quantitative readouts rather than secondary validations.

An alternative interpretation of the loss of BLI in FVB/N mice is that the progressive osteoblastic bone formation physically displaced or eliminated tumor cells from the bone marrow. This result is consistent with our data, including pathological analysis of the bones. Expanding trabecular architecture could progressively restrict the marrow space available for tumor cell occupation, thus reducing BLI signal without representing an immunologically targeted tumor elimination. The increased tissue depth of tumor cells within the mineralized matrix in osteoblastic lesions may also attenuate luminescent signal [25], contributing to the observed BLI decline independent of changes in tumor cell number or viability. However, R7 mammary carcinoma cells lost BLI signal after IT injection, yet produced no bone remodeling, demonstrating that osteoblastic bone formation alone is not responsible for the decline in luminescence.

Despite the loss of BLI signal, histological analysis of FVB/N bones at endpoint confirmed the presence of tumor cells within osteoblastic lesions, demonstrating that BLI signal loss of PyMT-CK(OB) tumors does not reflect complete tumor clearance. Together, these observations indicate that progressive BLI decline in PyMT-CK(OB) IT bones reflects a combination of immune-mediated reporter silencing, tissue attenuation through dense bone, and potentially reduced marrow space, rather than tumor elimination.

Intratibial injection provides a highly reproducible and experimentally tractable model of tumor-bone interaction. Both a pro and con of this injection site is that it bypasses critical steps of the metastatic cascade including intravasation, immune evasion in circulation, and homing to the bone, focusing the experiment on colonization [3, 14]. The surgical trauma associated with IT injection requires a localized wound-healing response that may amplify bone remodeling signals through pathways either active or not active during natural metastatic colonization [39]. This interpretation is supported by the absence of osteoblastic remodeling after intracardiac injection of PyMT-CK(OB) cells, despite confirmed skeletal dissemination, suggesting that the bone formation response is at least in part dependent on the inflammatory context created by direct intraosseous delivery. While PBS injection alone is not sufficient to drive abnormal bone formation, the wound healing response after injection could contribute to tumor colonization and osteoblastic remodeling at the metastatic site. Future studies could include sham IT injection with IC tumor inoculation to investigate this contribution. Alternatively, IT injection puts cancer cells directly in touch with one another at the metastatic site with neighboring cells. Pre-clustering of tumor cells prior to injection, shown to enhance metastatic efficiency and colonization in other models [40–42], represents an additional variable that could be tested in future work to determine whether cell contact in clusters shifts the osteoblastic/osteolytic balance observed here. Accordingly, conclusions drawn from IT injection models should be interpreted with awareness of the inflammatory and physical components. The aggressiveness of PyMT colonization also produced lesions too severe for standardized µCT analysis at later timepoints in some cohorts, suggesting that optimization of injection cell number and analysis timepoints is critical for maximizing the utility of this model in intervention studies.

The generation of PyMT-CF cells as a syngeneic osteolytic model from the same parental PyMT-CL background as of the osteoblastic PyMT-CK(OB) cells is a valuable experimental complement to PyMT-CK(OB). The availability of two cell lines – one osteoblastic, one osteolytic – derived from the same origin and injectable into the same immunocompetent syngeneic host enables controlled mechanistic comparisons of tumor-bone remodeling mechanisms without confounding variables introduced by different host backgrounds or immune status. The molecular basis for the divergent bone remodeling phenotypes between PyMT-CK(OB) and PyMT-CF despite shared parental origin is not yet established.

Interestingly, in vitro investigation of PyMT-CK(OB) and PyMT-CF cells on osteoblast mineralization reflected the mixed metastatic effect of PyMT-CF in these murine cells as secreted signaling from this line partially inhibited osteoblast mineralization. In contrast, osteoblast coculture with PyMT-CK(OB) cells did not result in increased mineralization compared to osteoblasts in monoculture, suggesting involvement of other stromal or marrow cells may contribute to the osteoblastic behavior of this cell line *in vivo*, which was further supported by increases in inflammatory cytokines in the PyMT-CK(OB) cocultures. The distinct protein secretion in PyMT-CK(OB) but not PyMT-CF cocultures compared to osteoblast monoculture reflects the osteoblastic nature of the former and suggests that PyMT-CF cells form metastatic lesions through communication with other cells, mostly likely osteoclasts. Ongoing characterization of secreted osteoclastogenic and osteoblastogenic cytokines may clarify the relevant upstream signals that drive osteoblastic versus osteolytic metastatic tumors.

The divergent responses of PyMT-CK(OB) and PyMT-CF tumors to zoledronate treatment further illustrate the functional distinction between these models. Zoledronate reduced PyMT-CK(OB) growth by BLI but did not prevent osteoblastic bone remodeling, consistent with a direct anti-tumor effect rather than a mechanism dependent on osteoclast suppression [27, 43]. This is unsurprising given that osteoblastic lesions are driven by excess bone formation and mineralization and aligns with clinical observations that bisphosphonates show limited efficacy in osteoblastic disease., as demonstrated by the failure of pamidronate and zoledronate to reduce skeletal related events in clinical trials of prostate cancer bone metastases [44, 45]. Indeed, despite demonstrated anti-tumor activity of bisphosphonates in preclinical models, the doses required are unlikely to be clinically achievable, and large clinical meta-analyses demonstrate benefit is confined to post-menopausal women, suggesting that bisphosphonate efficacy is largely microenvironment-dependent rather than tumor-directed [46–48]. In contrast, to PyMT-CK(OB), zoledronate significantly reduced both tumor growth and osteolytic remodeling in PyMT-CF bones, consistent with its established mechanism of osteoclast inhibition and supporting the interpretation that the mixed osteolytic PyMT-CF lesions are predominantly osteoclast-mediated. Together with the in vitro finding that PyMT-CF coculture suppresses osteoblast mineralization and ALP activity while PyMT-CK(OB) coculture does not, the zoledronate data provide further evidence that the sublines engage distinct bone remodeling programs. These complementary pharmacological and in vitro results support the distinction and functional relevance of these paired models for preclinical evaluation of bone targeted therapies.

Together, these results establish PyMT-CK(OB) intratibial injection in FVB/N mice as a characterized immunocompetent model of osteoblastic breast cancer bone metastasis and articulate a practical framework for interpreting bioluminescent and µCT data in this and related bone metastasis experiments. Future applications of this model include evaluation of immune-dependent bone remodeling, characterization of osteoblast and osteoclast activity by histomorphometry and serum marker analysis, manipulation of bone marrow cell populations through bone marrow transplantation and adoptive cell therapy, and preclinical testing of bone-targeted therapeutic strategies.

The complementary PyMT-CK(OB)/PyMT-CF syngeneic pair enables mechanistic dissection of osteoblastic and osteolytic bone metastasis biology in an immune-intact host.

## Methods

### Cell lines and culture

PNA-Luc-EZ-BoM is a luciferase- and GFP-expressing PyMT mammary carcinoma subline with high bone metastatic potential (gift from Dr. Siyuan Zhang, UT Southwestern) [22]. MMTV-PyMT-Luc (PyMT-CL) is a low-expression luciferase-positive mammary tumor cell line derived from a bone metastasis of an MMTV-PyMT mouse (gift from Dr. <u>C</u>onor <u>L</u>ynch, Moffitt Cancer Center) [20]. PyMT-CK(OB) (made by <u>C</u>ourtney <u>K</u>ing; <u>o</u>steo<u>b</u>lastic) and PyMT-CF (made by <u>C</u>ourtney <u>F</u>latt; heterogeneous/osteolytic) are lentiviral luciferase-expressing sublines of PyMT-CL generated in this work (described below). R7 cells are an estrogen receptor-positive murine mammary carcinoma line derived from MMTV-RON transgenic tumors [34, 35], and R7-luc cells are a lentiviral luciferase-expressing subline generated in this work (described below).

All murine PyMT-derived cell lines were maintained in Dulbecco’s Modified Eagle Medium (DMEM) supplemented with 10% fetal bovine serum (FBS) and 1% penicillin-streptomycin at 37°C and 5% CO_2_.

### Lentiviral transduction and cell line generation

For lentiviral packaging, HEK293T cells were transfected with the target plasmid and packaging constructs (PMDL, REV, VSVG) using calcium chloride transfection. Viral supernatant was collected at 48 and 72 hours post-transfection, concentrated using 5X Lentiviral Concentrator (Origene, Cat. #TR30026) and stored at -80°C.

To generate PyMT-CK(OB) and PyMT-CF cell lines, PyMT-CL parental cells were transduced with a lentiviral luciferase expression vector (Lenti-luciferase-P2A-Neo; Addgene plasmid #105621, MTA from Christopher Vakoc) containing a neomycin resistance selection cassette [49]. PyMT-CL cells were seeded at 4 × 10^4^ per well in 6-well plates and infected with 20-50 µL concentrated lentivirus per well in the presence of 8 µg/mL polybrene. After 24 hours, media was replaced, and cells were selected with 1000 µg/mL G418 disulfate for two to three weeks until uninfected control cells died from treatment. Stable luciferase expression was confirmed by bioluminescence assay. PyMT-CK(OB) and PyMT-CF represent independently transduced and selected sublines from the same parental PyMT-CL cells.

R7-luc cells were generated from R7 cells [34, 35] by the same lentiviral transduction protocol as described above.

### Bioluminescence assay

For in vitro bioluminescence quantification, cells were seeded in equal numbers in triplicate wells of 96-well plates and allowed to adhere for at least one hour. D-luciferin potassium salt (GoldBio, Cat. #LUCK-1g) was prepared as a 15 mg/mL stock in sterile DPBS and stored at -20°C. Cells were incubated with D-luciferin at 150 µg/mL final concentration and luminescence measured in a plate reader (SpectraMax iD5, Molecular Devices) after an empirically determined incubation period (0-20 minutes) sufficient to reach peak photon flux.

### Animals

All animal studies were conducted with approval from the University of Notre Dame Institutional Animal Care and Use Committee (protocols 21-11-6912, 21-12-6910, 21-12-6936, 21-12-6937, 24-10-8880, 24-10-8883, 24-10-8884) in accordance with guidelines from the National Institutes of Health and U.S. Department of Defense. Animals were maintained under specific pathogen-free conditions in the University of Notre Dame Freimann Life Sciences animal facility.

Female FVB/N mice (Charles River, #559NCIFVB) and NOD.CB17-*Prkdc^scid^*/NCrCrl (NOD SCID; Charles River) mice were used in this study.

### Cell preparation for in vivo injection

Cancer cells were harvested with 0.05% trypsin-EDTA at 80-90% confluency and resuspended in Dulbecco’s PBS with Ca^2+^ and Mg^2+^ (DPBS; Sigma-Aldrich, Cat. #D5652) for intratibial injection. For intracardiac injection, cells were resuspended in DPBS without Ca^2+^ or Mg^2+^. Cell viability and count were assessed by hemocytometer prior to injection.

### Intratibial injection

Mice were anesthetized with isoflurane and both hindlimbs shaved and sanitized with alcohol swabs. Cell suspensions of 1 × 10^4^ or 1 × 10^5^ cells in 10 µL DPBS were loaded into a 27-gauge needle (BD, Cat. #305109) coated with 2% bovine serum albumin in PBS to prevent cell adhesion. A second sterile 27-gauge needle was used to drill through the patellar ligament and tibial growth plate to access the medullary cavity, after which the cell-loaded needle was inserted and the suspension injected slowly. The procedure was repeated for the contralateral limb. Animals received ketoprofen (5 mg/kg) analgesia post-procedure and were monitored during recovery.

### Intracardiac injection

Mice were anesthetized with isoflurane and their chests were shaved and sanitized. The injection site was identified at the midpoint of the sternum, approximately 2 mm left of the midline. Cells (1 × 10^5^ in 100 µL DPBS without Ca^2+^/Mg^2+^) were loaded into a 29-gauge insulin syringe (Covidien, Cat. #8881511110) coated with 2% BSA in PBS, with a small air bubble to visualize arterial blood pulsation, confirming left ventricular placement. Cells were injected slowly between cardiac pulses. Successful injection was confirmed by immediate retroorbital D-luciferin injection and BLI showing a dilute, systemic bioluminescent signal. Mice with failed injections (intense focal chest signal) were euthanized and excluded from further analysis.

### In vivo bioluminescence imaging

Mice were imaged under isoflurane anesthesia in an IVIS Lumina II system (Caliper Life Sciences). D-luciferin (150 mg/kg body weight) was administered by intraperitoneal injection 15 minutes prior to imaging. Bioluminescence images were acquired using auto-exposure settings. Total photon flux (photons/second) was quantified by drawing regions of interest (ROIs) around both hindlimbs using Living Image software (version 4.2). Biweekly imaging was collected throughout the study duration.

### In vivo hypoxia imaging

To assess regional tumor hypoxia, the IVISense Hypoxia CA IV 680 Fluorescent Probe (HypoxiSense; PerkinElmer, Cat. #NEV11070) was administered via tail vein injection at 2 nmol per approximately 25 g body weight (100 µL). Fluorescence was imaged on the IVIS Lumina II at 24 and 48 hours post-injection. ROIs were drawn over the hindlimbs and the fluorescence signal was quantified with abdominal background subtraction. BLI was assessed simultaneously to compare the luciferase signal with hypoxia probe accumulation.

### Zoledronate treatment

Mice received intratibial injection of PyMT-CK(OB) or PyMT-CF tumor cells and were monitored by BLI to confirm successful engraftment. On the treatment start day (Day 3–8 post-injection, as indicated in individual experiments), mice were randomized by BLI tumor burden into balanced treatment cohorts. Zoledronate (Selleck Chemicals, Cat. #S1314; 5 μg/kg) or vehicle control were administered 3x/week s.c. for the duration of the study.

### Micro-computed tomography (µCT)

Following euthanasia and bone collection, tibiae were fixed overnight in 4% paraformaldehyde and stored in 70% ethanol. Bones were scanned on a Scanco Medical µCT-80 system (Scanco Medical, Brüttisellen, Switzerland) at 70 kVp, 114 µA, 200 ms integration time, and 10-µm isotropic resolution. Images were reconstructed with beam-hardening corrections and calibrated for bone mineral density. A Gaussian filter (variance = 1, support = 2) was applied for image smoothing, and a global threshold of 1324 mg HA/cm^3^ was used to segment mineralized tissue.

For trabecular analysis of IT injection tibiae, the region of interest (ROI) was defined in the proximal tibial metaphysis beginning at the growth plate and extending 1 mm distally into the medullary canal. The outer cortex was contoured and excluded from the trabecular ROI. Trabecular morphometric parameters were calculated using the Scanco trabecular morphology protocol, including bone volume fraction (BV/TV), trabecular number (Tb.N), trabecular thickness (Tb.Th), and trabecular separation (Tb.Sp). Bones were classified as osteoblastic or osteolytic based on Tb.N values exceeding or falling below two standard deviations from the mean of PBS-injected control bones in the same experiment, respectively. Bones with severe structural damage precluding reliable ROI definition were excluded from quantitative analysis and noted as degraded.

### Bone histology

Paraformaldehyde-fixed bones in tissue cassettes were transferred to 10% w/v EDTA solution (pH 7.6; EMD, Cat. #EX0534-1) and decalcified for 2-3 weeks at 4°C with agitation and weekly solution changes until mineral softening was confirmed by physical assessment. Decalcified bones and soft tissues were processed through an automated tissue processor (Leica TP1020), embedded in paraffin, and sectioned at 4-6 µm thickness.

Bone sections were deparaffinized in a 60°C oven for approximately one hour, rehydrated through three xylene washes and an ethanol gradient (100%, 95%, 70%), and rinsed in reverse osmosis (RO) water. Slides were stained with Hematoxylin 560 (Leica Biosystems, Cat. #3801570) for 4 minutes 30 seconds, differentiated in Define MX (Leica Biosystems, Cat. #3803595), blued in Blue Buffer (Leica Biosystems, Cat. #3802915), and counterstained with Eosin 515 Phloxine (Leica Biosystems, Cat. #3801606) for 1 minute 30 seconds. Slides were dehydrated, cleared in xylene, and mounted with CytoSeal XYL.

### Hydrogel culture

PyMT-CK(OB) and PyMT-CF cells were maintained as previously described. MC3T3-E1 murine osteoblastic cells (ATCC) were cultured in Minimum Essential Medium, Alpha modification (αMEM, Corning) and supplemented with 10% fetal bovine serum (FBS, Cytiva) and 1% Antibiotic Antimycotic Solution (AB/AM, Corning). Hydrogel fabrication using gelatin (Type A, 175 bloom, Sigma) crosslinked with transglutaminase (Moo Gloo Transglutaminase GS Formula) was completed as previously described [31]. Briefly, gelatin was mixed in complete αMEM media at a concentration of 900 mg/mL and transglutaminase was mixed with complete αMEM media at 0.3% per gram of gelatin (270 mg/mL). Nano-hydroxyapatite (nano-HA) particles were created on the day of experiment by mixing a 0.02 M calcium chloride dihydrate solution (2.92 mg/mL calcium chloride dihydrate in ultrapure water) with a phosphate solution (50mM HEPES buffer, 140 mM sodium chloride, 5 mM sodium hydroxide, 0.012 M sodium phosphate tribasic dodecahydrate, 0.017% v/v Darvan 821A in DI water) at a 1:2 ratio with DI water and sterile filtered using a 0.2 μm filter. Cells were suspended in media at a density of 2 × 10^6^ cells/mL and mixed 1:1 with the nano-HA solution prior to fabrication.

To fabricate hydrogels, the gelatin solution was heated to 37°C and mixed with the transglutaminase solution and cell suspension/nano-HA solution at a 1:1:1 ratio and pipetted into custom-made polydimethylsiloxane (PDMS) moulds. Osteoblasts were pipetted into 3 mm × 7 mm × 16 mm moulds. Cancer cells PyMT were pipetted into separate 3 mm × 4 mm × 13 mm moulds and allowed to crosslink at 4°C for 8 min. Hydrogels were left at room temperature for a minimum of 30 minutes before culturing at 37°C for the duration of the experiments. Hydrogels containing MC3T3 cells were cultured alone (MC3) or in the same well as hydrogels containing PyMT-CK(OB) (MC3+ PyMT-CK(OB)) or PyMT-CF (MC3+ PyMT-CF) cells for 28 days in osteogenic media (complete αMEM supplemented with 10 mM β-glycerophosphate, 100 nM dexamethasone, and 50 μM L-ascorbic acid).

### Biochemical analyses

Calcium content was quantified using a Calcium Liquicolor Kit (Stanbio Laboratories or Millipore Sigma) following manufacturer’s protocol. Briefly, hydrogels (N=3 or N=6) were removed from culture on days 7, 14, 21, or 28 and fixed for 24 hours at 4°C. Hydrogels were cut in half and digested in 1 mL of 1M hydrochloric acid (HCl) overnight at 60°C under constant rotation. Ten microliters of digested samples and standards were added to individual wells in a 96 well plate in triplicate followed by 200 μL of working solution. The plates were incubated for 10 min at room temperature before measuring absorbance at 550 nm on a plate reader (PerkinElmer LLC).

Extracellular ALP was measured in the cell culture medium (N=3) from days 7, 14, 21, and 28. Media was stored at -80°C before the assay was performed. A colorimetric assay of enzyme activity (SIGMAFAST p-NPP Kit) was used to determine ALP levels. P-nitrophenyl phosphate (pNPP) was used as the phosphate substrate with ALP enzyme as a standard. Cell culture media was added at 40 μL to a 96 well plate in triplicate with 50 μL of pNPP. The plate was incubated for 1 hour at room temperature in the dark before measuring absorbance at 405 nm on a plate reader.

TGF-β content was measured from cell culture medium using the Quantikine^TM^ ELISA Human/Mouse/Rat/Porcine/Canine TGF-β1 Immunoassay (Biotechne, R&D Systems, Cat. #DB100C) following the manufacturer’s protocol.

Protein expression of 111 mouse cytokines and chemokines was measured using the Proteome Profiler Mouse XL Cytokine Array (Biotechne, R&D Systems, Cat. #ARY028) from 1 mL cell culture medium stored at -80°C on day 7 from manufacturer’s protocol and imaged using the ChemiDoc (Biorad) for 3-12 minutes.

### Immunofluorescent staining

DAPI (1:2000; Millipore Sigma #D9564) and FITC (1:400; Fisher Scientific #50-225-0144) staining was performed on day 28 to analyze MC3T3 morphology. Hydrogels were permeabilized by incubating in 0.5% Triton-X in PBS for 10 min at 4°C. The gels were washed in BSA (Sigma A7906) three times and then incubated in BSA for 1 hour at room temperature. Constructs were incubated in phalloidin-FITC and DAPI dilactate for 30 minutes at room temperature and rinsed with 1% BSA solution. Constructs were stored in PBS at 4°C until fluorescence was detected using confocal laser scanning microscopy (Nikon AXR).

### Statistical analysis

Statistical analyses were calculated using GraphPad Prism (Version 10.0). Comparisons between two groups used Welch’s t-test or Mann-Whitney test, as indicated. Multi-group and multi-timepoint comparisons used two-way ANOVA with appropriate post-hoc tests for multiple comparisons. Statistical tests and significance thresholds are specified in individual figure legends. Data are presented as mean ± standard deviation unless otherwise noted.

## Acknowledgments

We would like to thank the members of the Littlepage and Niebur labs as well as Cristina Sanchez de Diego for their helpful insights and comments throughout this study. We thank the Friemann Life Sciences Center staff at the University of Notre Dame for the animal care and husbandry of mice used in this study. We also acknowledge the Notre Dame Integrated Imaging Facility (NDIIF) for imaging instrumentation and the Harper Cancer Research Institute and Tissue Core for instrumentation and support.

